# Evaluation of two Sigma Transwab^®^ systems for Maintenance of Viability of pathogenic *Candida* spp. Using the Clinical and Laboratory Standards Institute M40-A2 Standard

**DOI:** 10.1101/321018

**Authors:** E. Elcocks, E.C. Adukwu

## Abstract

A key aspect to routine microbiology processes include the retrieval, transport and maintenance of specimens. Swab transport systems (STS) can be utilised for their low cost, ease of use and their ability to recover and maintain specimens over long durations. An increase in healthcare complications due to fungal infections raises the requirement for STS to efficiently recover and preserve pathogens of yeast origin. The Clinical and Laboratory Standards Institute (CLSI) M40-A2 protocol is used to assess the compliance of STS to a quality control standard but at present does not include the recovery of yeast. The aim of this study was to compare the results of two commercial STS and their ability to recover and maintain viability of five clinical and reference strains of *Candida* spp., including *C. auris*, when stored at room temperature and 4°C, over 48 h, using the qualitative roll plate method. Findings from this study indicate that the STS used in this study are suitable for the collection and maintenance of the *Candida spp*. tested, and is very suitable for the recovery of clinical *C. auris*.

## INTRODUCTION

There has been a global increase of infections caused by invasive fungi, with molds and yeasts repeatedly described as pathogens (1). Of these infections, it is indicated that *Candida* spp. are a leading cause, and candidiasis effects more than a quarter of a million patients worldwide every year (2). This increase in opportunistic fungal pathogens can be linked to the progression of medical actions such as chemotherapy and transplants which results in an increase in immuno-compromised patients (3).

One species of *Candida* that is of particular concern is *C. auris*. Since the discovery of *C. auris* in 2009 (4) there have been rising concerns due to its rapid spread across the globe in the short time since its discovery (5); common misidentification for other *Candida* spp. (6–9); the serious associated infections including reports of isolation from the bloodstream, urinary tract, ear canal, wounds, heart muscle and bone (10); high mortality rate (10–12); and it’s antifungal resistance (6, 13). Rapid and efficient isolation and identification of these infectious agents are paramount to successful treatment, thus swab transport systems (STS) are often used for their low cost, ease of use and ability to maintain microorganism viability over extended periods of time (14). The Clinical and Laboratory Standards Institute (CLSI) M40-A2 is an approved standard which outlines testing procedures for liquid transport systems and provides manufacturers and end-users with a criteria for compliance (15). This protocol indicates the methodology to be used, storage conditions and length of incubation time for testing various STS. The protocol also outlines a list of organisms to be used for quality control, though amongst these organisms, yeasts are only indicated for urine transport systems. The compliance criteria of an STS in M40-A2 is described as no greater than 3 log decrease when stored at 4 °C or room temperature, or 1 log increase when stored at 4 °C. The enumeration should also be ≥5 CFU from the same dilution as used in time-zero plate counts after the specified holding period. The aim of this study is to investigate and evaluate the efficiency of two commercial transport swabs to recover several *Candida* spp., including *C. auris*, using the qualitative roll-plate method outlined by the M40-A2.

## METHODS

The method in this study followed the CLSI M40-A2 protocol with some minor adaptations. The STS used in this study were both manufactured by Medical Wire and Equipment (MWE; Corsham, UK) and included their Sigma Transwab^®^ (MW176S) and Sigma Transwab^®^ Purflock^®^ (MW176PF). Both STS utilise a liquid amies carrying media. The Gram stain procedure was carried out on all species of *Candida. Candida* spp. stored on microbank beads at −80°C were grown on Sabouraud Dextrose Agar (SDA; EO Labs, Bonnybridge, UK) at 30°C for 48 h and used to inoculate 0.85% physiological saline (OXOID, Basingstoke, UK) until turbidity reached McFarland Standard 0.5 (OD_625nm_ 0.08–0.1). Serial dilutions were made in saline and 100 µl of 10^-2^ and 10^-3^ dilutions were used to inoculate swabs aiming to achieve an approximate inoculation spike of 10^5^ and 10^6^ CFU/mL. Initial inocula was verified by plate count to determine CFU/ml. Three swabs were used per condition and incubated at either room temperature or 4°C for 0, 24 and 48 hours (T0/T24/T48). Swabs were plated using the roll plate method outlined in M40-A2 onto SDA agar and incubated at 30°C for 48 h and then enumerated.

## RESULTS

Results from Gram staining *Candida* spp. can be seen in Figure 1. All show the expected Gram positive appearance with typical round/oval budding morphology. All 100 strains of *Candida*, including *C. auris*, were successfully recovered by both STS. Both STS used were compliant with the M40-A2 criteria for all *Candida* spp. tested (Table 1). No overgrowth of organisms is shown when stored at 4°C with most of the STS 103 showing a slight decrease in CFU after 48 h, whilst all STS show an increase in CFU after 48 h when stored at room temp (Figure 2).

**Figure 1.**
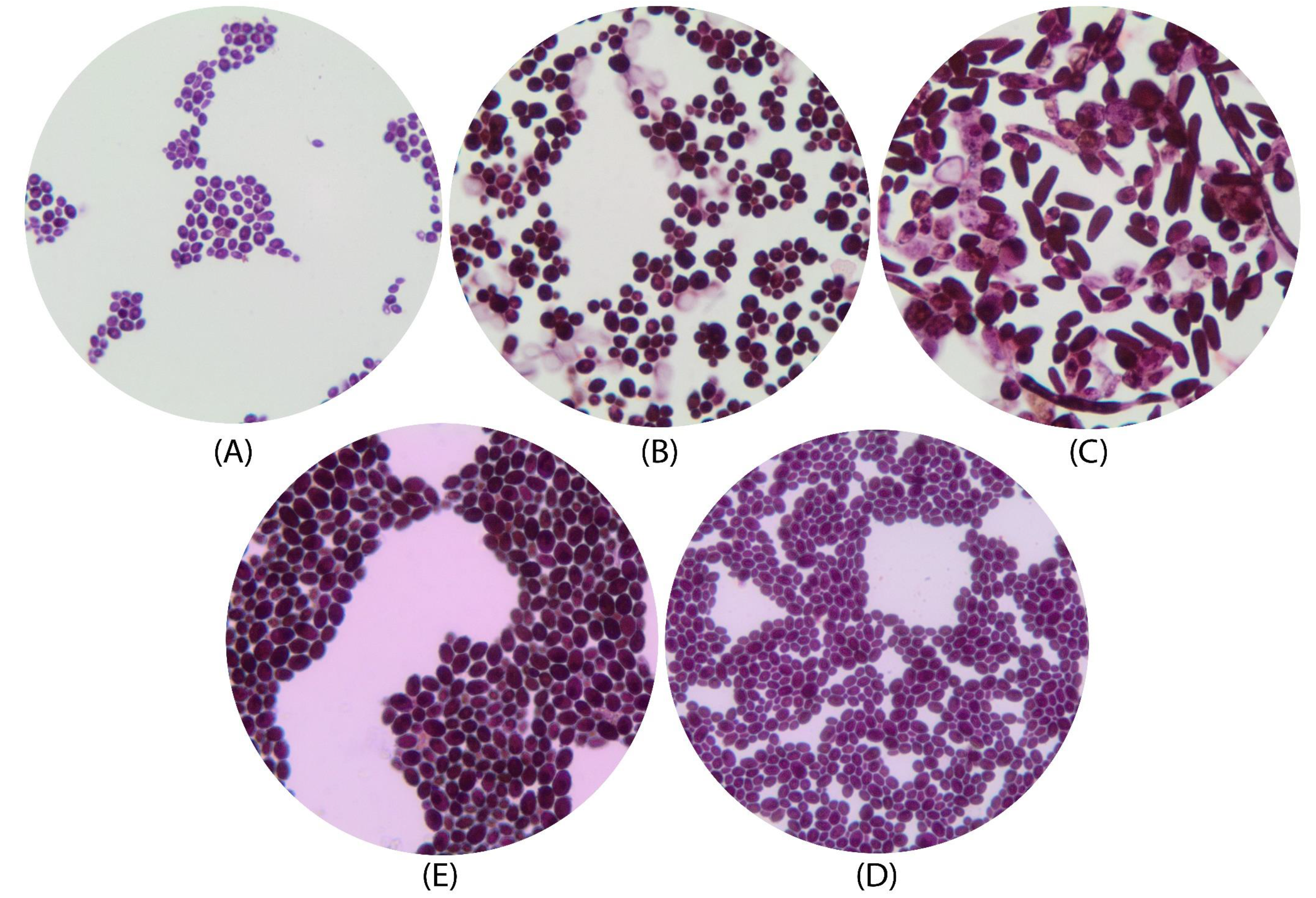
Gram strain images of: (A) *C. auris* NCPF 8971; (B) *C. albicans* NCPF 3179; (C) *C. tropicalis* NCPF 3111; (D) *C. glabrata* ATCC 2001; (E) *C. parapsilosis* ATCC 22019. 100x magnification.

**Figure 2.**
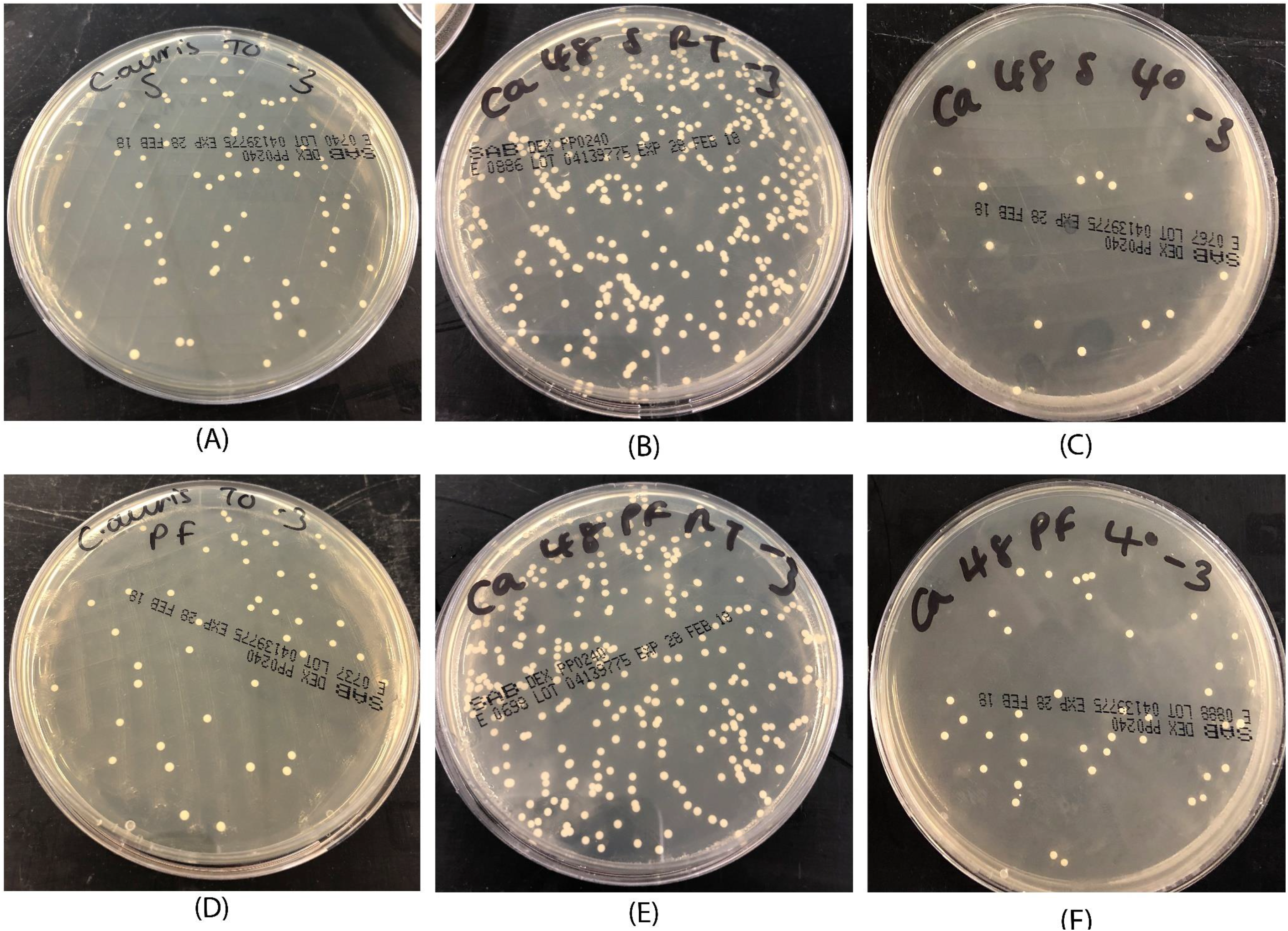
Enumeration plates showing growth of *Candida auris* on SDA for (A) Sigma STS at T0; (B) Sigma STS at T48 and RT; (C) Sigma STS at T48 and 4°C; (D) Sigma Purflock STS at T0; (E) Sigma Purflock STS at T48 and RT; (F) Sigma Purflock STS at T48 and 4°C.

**Table 1.** Qualitative results of bacterial recovery of *Candida* spp. isolated from the Sigma Transwab^®^ (MW176S) and Sigma Transwab^®^ Purflock^®^ (MW176PF) at room temperature and at 4°C.

| Organism | Swab | Temp. | Dilution 10<sup>-2</sup> |  |  |  | Dilution 10<sup>-3</sup> |  |  |  |
| --- | --- | --- | --- | --- | --- | --- | --- | --- | --- | --- |
|  |  |  | 0hr | 24hr | 48hr | Compliance | 0hr | 24hr | 48hr | Compliance |
| *Candida auris*<br>NCPF 8971 | Sigma | RT | TNTC | TNTC | TNTC | ✓ | 51 | 50 | TNTC | ✓ |
|  | Sigma | 4°C | TNTC | 179 | 183 | ✓ | 51 | 16 | 20 | ✓ |
|  | Purflock | RT | TNTC | TNTC | TNTC | ✓ | 35 | 55 | TNTC | ✓ |
|  | Purflock | 4°C | TNTC | TNTC | 208 | ✓ | 35 | 43 | 29 | ✓ |
| *Candida albicans*<br>NCPF 3179 | Sigma | RT | 164 | TNTC | TNTC | ✓ | 14 | 117 | 255 | ✓ |
|  | Sigma | 4°C | 164 | 153 | 134 | ✓ | 14 | 14 | 13 | ✓ |
|  | Purflock | RT | 176 | TNTC | TNTC | ✓ | 14 | 92 | 200 | ✓ |
|  | Purflock | 4°C | 176 | 100 | 133 | ✓ | 14 | 12 | 17 | ✓ |
| *Candida tropicalis*<br>NCPF 3111 | Sigma | RT | 128 | TNTC | TNTC | ✓ | 14 | 143 | TNTC | ✓ |
|  | Sigma | 4°C | 128 | 71 | 66 | ✓ | 14 | 4 | 5 | ✓ |
|  | Purflock | RT | 102 | TNTC | TNTC | ✓ | 12 | 67 | 146 | ✓ |
|  | Purflock | 4°C | 102 | 77 | 55 | ✓ | 12 | 5 | 9 | ✓ |
| *Candida parapsilosis*<br>ATCC 22019 | Sigma | RT | 158 | TNTC | TNTC | ✓ | 7 | 46 | 236 | ✓ |
|  | Sigma | 4°C | 158 | 58 | 109 | ✓ | 7 | 10 | 6 | ✓ |
|  | Purflock | RT | 135 | TNTC | TNTC | ✓ | 13 | 46 | 121 | ✓ |
|  | Purflock | 4°C | 135 | 120 | 154 | ✓ | 13 | 9 | 13 | ✓ |
| *Candida glabrata*<br>ATCC 2001 | Sigma | RT | 198 | 96 | 269 | ✓ | 16 | 8 | 20 | ✓ |
|  | Sigma | 4°C | 198 | 71 | 113 | ✓ | 16 | 4 | 10 | ✓ |
|  | Purflock | RT | 171 | 182 | 260 | ✓ | 18 | 21 | 36 | ✓ |
|  | Purflock | 4°C | 171 | 197 | 165 | ✓ | 18 | 17 | 23 | ✓ |

TNTC – too numerous to count; RT – Room temperature. (n=3)

## DISCUSSION

Both swab transport systems have demonstrated successful and efficient recovery 108 and maintenance of *Candida* spp. and of particular interest is the recovery of *C. auris*. Considering the recent increase in global concern for *C. auris* and its associated implications, as previously mentioned, and in light of the findings from this study, MWE Sigma Transwab^®^ and Sigma Transwab^®^ Purflock^®^ can be described as very suitable for the recovery of *C. auris* and can be implemented in the viability and transport of this opportunistic pathogen and aid in the timely treatment of associated infections. In a recent study (16) several Candida spp. were recovered using other commercial STS based on the M40-A2 protocol, however, the study did not investigate three of the Candida spp. in this study including the *C. auris*.

To our knowledge, this study is the first to evaluate STS efficiency and recovery of *C. auris* using the M40-A2 protocol. The M40-A2 does not currently directly address the recovery of yeasts, with the exception of guidance for urine transport systems (15).

In this study it was found that when using the protocol for adjusting initial inocula, enumeration was lower than that of bacteria, as expected, due to the difference in size of yeast and bacterial cells. Of the *Candida* spp. used within this study, only *C. auris* is a known clinical isolate. Though using a panel of clinically isolated bacteria would better reflect the use of STS in clinical situations, the use of reference strain cultures is in-line with the M40-A2 protocol which utilises quality control strains for testing STS. With regards to a direct comparison of each STS, though the swab tip of each STS used in this study varied, the results from the foam or flocked swab were similar. One study (17) claims that foam swabs are superior to flocked swabs, though this preference was due to foam swabs ability to perform better when used in antigen testing experiments and not a reflection of the specimen recovery and maintenance.

Within this study, overgrowth was seen in both swabs for *C. auris* and in the Sigma swab for *C. albicans* at room temperature. When stored at 4°C, no overgrowth was seen in either STS. With an increase of transport delays being seen due to increasing costs for containment and movement of services to more central laboratories^[18]^, transport in cool temperatures is suitable, particularly for specimens of pathogenic yeast origin, and specimen maintenance would be ensured.

## ACKNOWLEDGEMENTS

This work was funded by Medical Wire and Equipment, Corsham, UK and supported by the Faculty of Health and Applied Sciences, University of the West of England, Bristol, UK. The funders had no role in data collection and interpretation, or the decisions involved with submitting the work for publication.

